# PDS5s control the *Arabidopsis* 3D genome by suppressing the formation of TAD-like domains

**DOI:** 10.1101/2024.07.31.606105

**Authors:** Anna-Maria Göbel, Sida Zhou, Zhidan Wang, Sofia Tzourtzou, Axel Himmelbach, Shiwei Zheng, Mónica Pradillo, Chang Liu, Hua Jiang

## Abstract

One key structural element in the three-dimensional (3D) chromatin organization is the Topologically Associating Domain (TAD), which facilitates the formation of distinct chromatin compartments and fosters specific chromatin interactions. Similar chromatin compartments, known as TAD-like domains, have been identified in many plant species. However, the model plant *Arabidopsis* thaliana has been an exception. In this study, we address this long-standing issue by presenting evidence that *Arabidopsis* PDS5 proteins play a crucial role in preventing the formation of TAD-like domains.

---

Being structurally analogous to TADs, TAD-like domains are higher-order chromatin structures in which interactions between DNA loci within the TADs are enhanced and the boundaries of the TADs show chromatin insulation^1,2^. The absence of prominent TAD-like structures in the *Arabidopsis* genome, which were otherwise found in many plant species, has puzzled the plant science community for over ten years^3-5^. In animals, the cohesin complex, regulated by the insulator protein CTCF, plays a pivotal role in defining TAD borders where chromatin insulation takes place^2,6^. Assuming functional conservation of cohesin in plants and animals^7^, we initiated some trial Hi-C experiments to assess the potential impact of the perturbation of *Arabidopsis* cohesin activities on chromatin organization (Supplemental Data 1). Because loss-of-function mutants of *Arabidopsis* cohesin subunits SMC1 and SMC3 are not viable, we instead selected mutants that likely influence cohesin-chromatin interactions^8^. To this end, *wapl1 wapl2* double mutants (hereafter referred to as *wapl1/2*) and *pds5a pds5b pds5c pds5e* quadruple mutants (hereafter referred to as *pds5a/b/c/e*) were used^9-11^.

On a chromosomal level, *wapl1/2, pds5a/b/c/e*, and wild-type (WT) Hi-C maps appeared similar (Supplemental Fig. 1a and Supplemental Fig. 2). However, when we zoomed in on the diagonal line to examine details in local chromatin contacts (i.e., contacts that span distances up to a few hundred kilo base-pairs), remarkable differences in the *pds5a/b/c/e* plants were observed (Fig. 1a, Supplemental Fig. 1a-c). In *pds5a/b/c/e*, conspicuous TAD-like domains were ectopically formed and widely distributed through the genome except for the pericentromeric regions which were constitutive heterochromatin. Concordantly, chromatin interactions decay plots, which revealed how chromatin contact strength decayed as a function of genomic distance, showed specific changes in chromosome arm regions in *pds5a/b/c/e* in comparison to the WT sample, indicating that the former adopted distinct chromatin packing patterns (Supplemental Fig. 1d).

**Figure 1.**
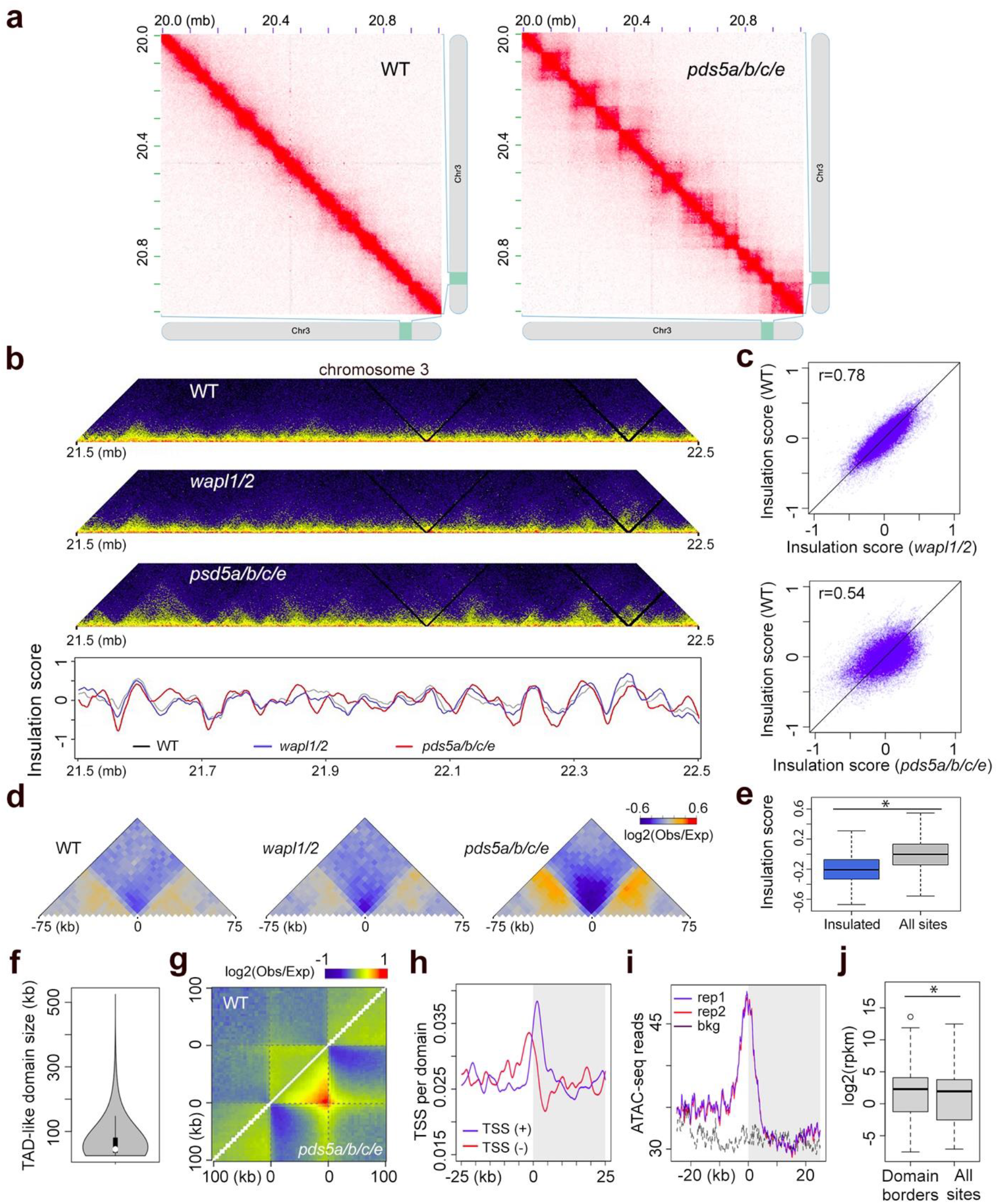
Analysis of TAD-like domains in *pds5a/b/c/e*. (a) Prominent TAD-like structures in *pds5a/b/c/e*, showcased by a representative 1 Mb genomic region. (b) Comparison of insulation score profile of a 1 Mb region at chromosome 3 with the corresponding Hi-C map shown on top. The regions showing local insulation score minima indicate strong chromatin insulation. (c) Genome-wide correlation of insulation scores between different plants. r, Pearson correlation coefficients. (e) Metagene plots of chromatin interaction patterns in different plants around the loci displaying strong chromatin insulation in *pds5a/b/c/e*. (d) Insulation scores in wild-type plants. The category “Insulated” refers to those loci showing strong chromatin insulation in *pds5a/b/c/e*. *, significant (p < 10^−16^) according to a two-sided Mann–Whitney U-test. (f) Size distribution of TAD-like domains in *pds5a/b/c/e*. The median domain size is 45 kb. (g) Metagene plot illustrates relative chromatin contact strengths in TAD-like domains identified in *pds5a/b/c/e*. The flanking regions (100 kb) are also included. The top side shows WT and the bottom side shows mutant. Besides the strong insulation region, the extended signal on the top of the triangle promotes the detection of strong boundaries in *pds5a/b/c/e*. (h) Distribution of TSS around *pds5a/b/c/e* TAD borders (labeled with 0). The right grey side represents the 25 kb within the TAD-like domains, the left side from 0 represents 25 kb of flanking area. Both transcription directions are included. (i) Chromatin accessibility associated with TAD-like domain borders in *pds5a/b/c/e*. (j) Comparison of gene expression in *pds5a/b/c/e* between genes located at TAD-like domain borders and elsewhere. *, significant (p < 10^−10^) according to a two-sided Mann–Whitney U-test.

The *Arabidopsis* genome encodes five homologs of *PDS5*, which are redundantly involved in not only chromatid cohesion and DNA damage repair but also small interfering RNA production and transcriptional regulation^12-15^. At a transcriptional level, *PDS5A* is the most actively expressed *PDS5* gene; furthermore, *PDS5A* was expressed approximately 6 times as its closest homolog, *PDS5B* (Supplemental Data 2). Compared to other PDS5 proteins, PDS5A and PDS5B have considerably more Armadillo-type fold domain regions, which potentially promote their interactions with proteins and nucleic acids^13^. To investigate the potential functional redundancy of *Arabidopsis PDS5* genes in regulating chromatin organization, we conducted further Hi-C experiments to examine single, double, and triple *pds5* mutants. For all those examined mutants that contained a *pds5a* allele, their genomes exhibited similar TAD-like domains as in the *pds5a/b/c/e*, indicating that *PDS5A* is the major factor suppressing TAD-like structures (Supplemental Fig. 3a). Supporting this notion, transforming *pds5a/b* with a construct bearing *PDS5A* genomic DNA rescued its chromatin organization patterns (Supplemental Fig. 3b-d). Given that *PDS5s* redundantly regulate *Arabidopsis* seedling development^13^ and that *pds5a/b/c/e* had the most altered transcriptome profile (Supplemental Fig. 4 and Supplemental Data 2), we presumed that chromatin organization in *pds5a/b/c/e* was affected the most compared to other lower-order mutants. Thus, *pds5a/b/c/e* Hi-C libraries were deep sequenced for further analyses (Supplemental Data 1).

Compared to WT, about 15% of *pds5a/b/c/e* chromatin showed AB Compartment swaps, reflecting mild rewiring of genome topology at a chromosomal level (Supplemental Fig. 5a-d and Supplemental Data 3). Interestingly, these AB compartment swaps were not linked to changes in active (e.g., H3K4me3) or repressive (e.g., H3K27me3 and H3K9me2) histone marks, which were enriched in A and B Compartment regions, respectively (Supplemental Fig. 5b-f and Supplemental Data 4). Furthermore, although A and B Compartment identities are generally linked to active and inactive gene expression, respectively, in *pds5a/b/c/e*, the switch in compartment identity was independent of changes in the transcriptome (Supplemental Fig. 5g, h).

Next, we inspected newly emerged local chromatin contact patterns on the *pds5a/b/c/e* Hi-C map. By calculating insulation scores, we identified genomic regions in *pds5a/b/c/e* showing strong chromatin insulation (Fig. 1b and Supplemental Data 5). Interestingly, most insulated regions in *pds5a/b/c/e* already exhibited weak insulation patterns in WT plants (Fig. 1c-e). Consistent with a recent report, these insulated regions were enriched with 5’ flanking regions of genes’ transcription start sites^16^ (Supplemental Fig. 6). TAD-like domains in *pds5a/b/c/e* were also annotated (Fig. 1f and Supplemental Data 6). Interestingly, these regions already showed weak domain formation propensity on the wild-type Hi-C map (Fig. 1g). Like other plant species, *pds5a/b/c/e* TAD-like domain borders were enriched with promoter regions, higher chromatin accessibility, and genes with higher expression levels (Fig. 1h-j and Supplemental Figure 7a), suggesting that these emerged TAD-like domain structures are linked to gene expression^17^. However, further examination of transcriptional activities in these regions, such as euchromatin mark deposition (H3K4me3), chromatin accessibility, RNA-polymerase II association, and gene expression levels, did not reveal noticeable changes in *pds5a/b/c/e* (Supplemental Fig. 8). Therefore, changes in chromatin insulation in *pds5a/b/c/e* did not cause a direct impact on gene expression.

Along with the emergence of TAD-like domains, the extensive changes in local chromatin organization in *pds5a/b/c/e* created a large number of new chromatin contacts. To better understand the chromatin interaction network in *pds5a/b/c/e*, we identified chromatin contacts with statistical significance (hereafter referred to as “chromatin loops”) in WT and mutant plants, respectively^18^. These two genotypes had similar chromatin loops in a short distance range (<10 kb), but the fraction of shared chromatin loops dropped dramatically at longer distances (10-100kb) (Fig. 2a). Among the long-distance chromatin loops (10-100kb), we found that about 40% of loop-forming regions in WT plants overlapped with protein-coding genes. In contrast, the percentage in *pds5a/b/c/e* was 65%, which was partially due to having fewer transposons overlapping with *pds5a/b/c/e* loops (Fig. 2a-c).

**Figure 2.**
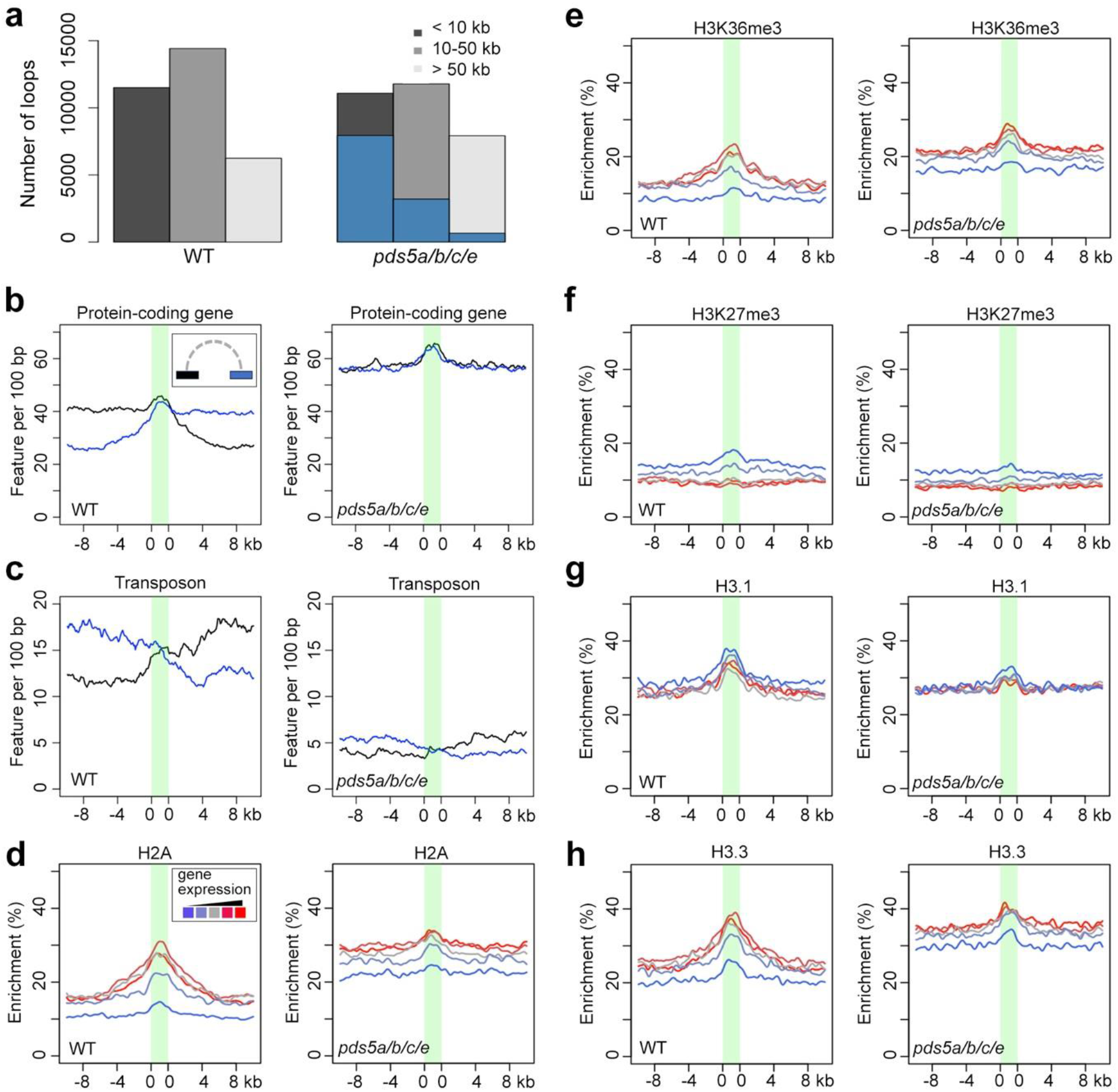
Comparison of epigenetic features associated with chromatin loops and gene expression between WT and *pds5a/b/c/e*. (a) Proportion of chromatin loop sizes in WT and *pds5a/b/c/e*. Loops smaller than 10 kb, between 10 and 50 kb, and bigger than 50 kb are included. Blue scaled bars indicate those loops in *pds5a/b/c/e* which are also identified in wild-type plants. (b, c) Genomic features at regions annotated with chromatin loops, which are illustrated as the green area in each plot. The blue and black curves refer to individual loop anchor regions. (d - h) Enrichment of epigenetic marks around chromatin regions that form loops with protein-coding genes. These regions are divided into five groups according to expression levels of their interacting protein-coding genes. The color gradient from blue via grey to red represents ascending gene expression.

Among the long-distance chromatin loops in *pds5a/b/c/e*, we examined their epigenetic profiles in relation to gene expression. At the moment, comprehensive datasets that depict various epigenetic marks in *pds5a/b/c/e* are not available. However, given that the WT and *pds5a/b/c/e* plants exhibited highly similar patterns of three representative epigenetic marks (i.e., H3K4me3, H3K9me2, and H3K27me3), and that these plants had comparable transcriptome profiles (Supplemental Figs. 7b and 9), we expected that at a genome-wide level, the epigenomic landscape of *pds5a/b/c/e* is similar to that of WT plants. With such an assumption, we analyzed several histone marks at loci forming chromatin loops with protein-coding genes, in which WT seedlings’ epigenetic features were used to approximate those in *pds5a/b/c/e*. We found that in both WT and *pds5a/b/c/e* plants, inactive genes tended to form chromatin loops with regions carrying repressive chromatin marks (e.g., H3K27me3), while actively expressed genes preferentially interacted with active chromatin regions (e.g., those decorated with H3K36me3 and H2A) (Fig. 2d-h). These results suggest that *pds5a/b/c/e* and WT chromatin reconstituted long-range chromatin contact networks with the same chromatin segregation principle, in which active and repressed chromatin form partners with similar epigenetic profiles, respectively.

In conclusion, our work reveals that PDS5A, probably in conjunction with other PDS5 homologs, largely suppresses long-range chromatin contacts and chromatin insulation, providing a genetic explanation for why *Arabidopsis* lacks distal enhancers as opposed to other plants. Although we did not extensively profile individual epigenetic marks, the emergence of TADs-like domains in *pds5a/b/c/e* appears independent from them, as three distinct types of histone marks (i.e., H3K4me3, H3K9me2, and H3K27me3) remained largely unchanged in the mutants. Future work would be oriented to understand why and how TAD-like chromatin structures become suppressed in *Arabidopsis*. Notably, PDS5A contains a Tudor domain, which shares a high degree of sequence and structural conservation with that of PDS5C (Supplemental Fig. 10). The PDS5C Tudor domain is known to recognize mono-methylated histone H3K4, suggesting a potential mechanism whereby PDS5A interacts with chromatin via its Tudor domain^15,19^. Besides, the identification of PDS5A as a major player in shaping the *Arabidopsis* genome topology offers valuable insights for discovering other structural proteins involved in plant 3D chromatin organization.

## Methods

### Plant materials and growth conditions

The *Arabidopsis thaliana* accession *Columbia* was used as wild-type (WT) reference. The *pds5a, pds5b, pds5c, pds5e, pds5a/b, pds5a/b/c, pds5a/b/c/e* and *wapl1/2* mutants were previously reported^13,20^. Seeds were treated at 4°C for 3 days before sowing on the soil. All plants were cultured in the glasshouse with a 16 h : 8 h, light : dark photoperiod at 22 °C.

### Plasmid construction

To make the *PDS5A:2HA* construct, a 10 kb genomic DNA containing the RNA-coding region of *PDS5A* was amplified as two overlapping fragments: oligos 5’-CTTTGTACAAGAAAGCTGGGTGACGTCGGTATTCTTACACTAATG -3’ and 5’-AGGGTATCCAGCATAATCTGGTACGTCGTATGGGTATCCTATTGCTGTCCTCGAGATTGA - 3 for the first fragment, while oligos 5’-CTTTGTACAAGAAAGCTGGGTCAAGAATAAGTAGTTGACATCA -3’ and 5’-GATTATGCTGGATACCCTTACGACGTACCAGATTACGCTTAGGTTTGCCGTATGCCGTA -3’ for the second fragment. The PCR products were cloned into the pFK206 vector^21^ via a Gibson Assembly reaction (NEB). The construct sequence was verified with Sanger sequencing and transferred into plants with the Agrobacteria-mediated floral dip method.

### *In situ* Hi-C and data analyses

In situ Hi-C libraries were generated following established protocols^22,23^. Each sample was subjected to two Hi-C library replicates, with approximately 0.5 g of fixed sample homogenized for nuclei isolation. The isolated nuclei were resuspended in 150 µl of 0.5% SDS and divided into three tubes. After a 5-minute incubation at 62°C for penetration, SDS was neutralized by adding 145 µl water and 25 µl 10% Triton X-100, followed by a 15-minute incubation at 37°C. Subsequently, chromatin was digested using 50 U Dpn II (NEB) in each tube overnight at 37°C. The following day, Dpn II was inactivated by a 20-minute incubation at 62°C. Sticky ends were then filled in by adding 1 µl of 10 mM dTTP, 1 µl of 10 mM dATP, 1 µl of 10 mM dGTP, 10 µl of 1 mM biotin-14-dCTP, 29 µl water, and 40 U Klenow fragment (Thermo Fisher), with an incubation at 37°C for 2 hours. Following the addition of 663 µl water, 120 µl of 10x blunt-end ligation buffer (containing 300 mM Tris-HCl, 100 mM MgCl2, 100 mM DTT, 1 mM ATP, pH 7.8), and 40 U of T4 DNA ligase (Thermo Fisher), proximity ligation was conducted at room temperature for a duration of 4 hours. Subsequently, the ligation products from the three tubes of identical samples were combined and resuspended in 650 µl SDS buffer (50 mM Tris-HCl, 1% SDS, 10 mM EDTA, pH 8.0). Following treatment with 10 µg proteinase K (Thermo Fisher) at 55°C for 30 minutes, de-crosslinking was performed by adding 30 μl 5 M NaCl and incubating overnight at 65°C. The DNA was recovered and treated with RNase A at 37°C for 30 minutes. After purification, 3∼5 μg of the recovered Hi-C DNA was adjusted to 130 μl with TE buffer (10 mM Tris-HCl, 1 mM EDTA, pH 8.0), and sheared using a Q800R3 (QSONICA) sonicator with the following settings: 25% amplitude, 15s ON, 15s OFF, pulse-on time for 4.5 min, to achieve a fragment size shorter than 500 bp. The sonicated DNA was purified with Ampure beads to recover fragments longer than 300 bp. Subsequently, in a 50 μl reaction volume, the DNA was mixed with 0.5 μl 10 mM dTTP, 0.5 μl 10 mM dATP, and 5 U T4 DNA polymerase, and incubated at 20°C for 30 minutes to remove biotin from unligated DNA ends. Following purification with Ampure beads, the DNA underwent end-repair and adaptor ligation using the NEBNext® Ultra™ II DNA Library Prep Kit (NEB). Following affinity purification using Dynabeads MyOne Streptavidin C1 beads (Invitrogen), the ligated DNA was subsequently subjected to amplification through 12 PCR cycles. The resulting libraries were then sequenced using an Illumina Novaseq instrument, generating reads with a length of 2 × 150 bp.

Reads mapping to the TAIR10 genome with Bowtie 2 (v2.2.4), removal of PCR duplicates, and reads filtering were performed as described. Hi-C reads of each sample are summarized in Supplemental Data 1. Hi-C map normalization was performed by using an iterative matrix correction function in the “HiTC” package in R programm. For all Hi-C maps, the iterative normalization process was stopped when the eps value, which reflected how similar the matrices in two consecutive correction steps were, dropped below 1 × 10^−4^. In addition, the filtered Hi-C reads were used to create *hic* files with the juicer tool for interactive Hi-C map inspection. The annotation of A/B Compartment of chromosome arms is available in Supplemental Data 3.

Insulation scores were calculated according to a previous study^24^. Here we used Hi-C maps normalized with 2 kb bins, and intrachromosomal contacts up to 50 kb were included in the computation. Because of chromosome translocation in the *pds5a/b/c/e* plants, genomic regions at the left arm of chromosome 1 and the left arm of chromosome 5 were not included. Subsequently, potential “insulated regions” were identified by looking for regions with local insulation score minima. This is done by implementing the *peak* function from the splus2R package in R with *span=10*. In this study, we annotated regions with insulation scores lower than -0.25 and overlapping with insulation score local minima as “insulated regions”. A list containing insulation scores and insulated region annotation is available in Supplemental Data 5.

TAD-like domains that emerged on the *pds5a/b/c/e* Hi-C map were identified with the “arrowhead” algorithm, which we have previously applied to similar chromatin features in the rice and Marchantia genomes^22,25,26^. In this study, we set the cutoff for the TAD score matrix as 0.95, the minimum number of filtered pixels belonging to a potential TAD as 6, and the minimum TAD score of pixels at TAD borders as 1.05. TAD annotation is available in Supplemental Data 6. The TAD calling pipeline was not applied to pericentromeric regions because they did not show clear changes on the Hi-C map (Supplemental Fig. 1). Furthermore, because of the translocation between chromosome 1 left arm and chromosome 5 left arm, these two chromosome arms were not subject to TAD calling (Supplemental Fig. 2).

Chromatin contacts with statistical significance were identified with the FiTHiC2 tool^18^. The resolution was set to be 2 kb, and the distance range in which the chromatin interaction model was built and chromatin loops were called was between 4 kb and 100 kb. After running FitHiC2, chromatin contacts with q-values less than 0.05 were used for downstream pattern analyses. The entire list of chromatin loop list is available in the figshare repository with the following link: https://figshare.com/s/8a653d26c484048b67bf

### Chromatin immunoprecipitation and ChIP-seq data analyses

Fourteen-day-old *Arabidopsis* leaves were fixed under vacuum for 30 minutes with 1% formaldehyde solution in MC buffer (10 mM potassium phosphate, pH 7.0, 50 mM NaCl, and 0.1 M sucrose) at room temperature. The fixation process was stopped by replacing the solution with 0.15 M glycine in MC buffer under vacuum for 10 minutes at room temperature. Approximately 0.5 gram of the fixed tissue was homogenized and suspended in nuclei isolation buffer (containing 20 mM HEPES, pH 8.0, 250 mM sucrose, 1 mM MgCl2, 5 mM KCl, 40% glycerol, 0.25% Triton X-100, 0.1 mM PMSF, and 0.1% 2-mercaptoethanol) and then filtered using double-layered miracloth (Millipore). The isolated nuclei were resuspended in 0.5 mL of sonication buffer (10 mM potassium phosphate, pH 7.0, 0.1 mM NaCl, 0.5% sarkosyl, and 10 mM EDTA). Chromatin was fragmented by sonication using a QSONICA sonicator Q800R3 to achieve an average fragment size of approximately 400 base pairs.

Next, 50 µl of 10% Triton X-100 was combined with the sonicated sample, and 25 µl of this mixture was retained as an input sample. The remaining sheared chromatin was mixed with an equal volume of IP buffer (comprising 50 mM HEPES, pH 7.5, 150 mM NaCl, 5 mM MgCl2, 10 µM ZnSO4, 1% Triton X-100, and 0.05% SDS) and incubated with antibodies (anti-Pol2, Abcam ab5408; anti-H3, Sigma H9289; anti-H3K4me3, Abcam ab8580; anti-H3K9me2, Diagenode C15410060; anti-H3K27me3, Millipore 07-449) at 4°C for 2 hours, followed by further incubation with protein A/G magnetic beads (Thermo Fisher) 4°C for 2 hours. The beads were subsequently washed at 4°C as follows: 2 washes with IP buffer, 1 wash with IP buffer containing 500 mM NaCl, and 1 wash with LiCl buffer (0.25 M LiCl, 1% NP-40, 1% deoxycholate, 1 mM EDTA, and 10 mM Tris-HCl pH 8.0), each for 3 minutes. After a brief rinse with TE buffer (comprising 10 mM Tris-HCl, pH 8.0, and 1 mM EDTA), the beads were resuspended in 200 µl of elution buffer (containing 50 mM Tris-HCl, pH 8.0, 200 mM NaCl, 1% SDS, and 10 mM EDTA) and incubated at 65°C for 6 hours. This was followed by Proteinase K treatment at 45°C for 1 hour. DNA was purified using the MinElute PCR purification kit (QIAGEN) and subsequently converted into sequencing libraries using the NEBNext® Ultra™ II DNA Library Prep Kit (NEB).

After sequencing, ChIP-seq reads were aligned to the TAIR10 genome using Bowtie 2 (version 2.2.4). The mapped reads were used to generate coverage files with 100 bp windows using bedtools (v2.26.0). The mapped reads were analyzed by MACS2 v2.2.7.1. The “--broad” flag was on during peak calling for H3K27me3. The reads from the anti-H3 sample used as control and the other parameters are -q 0.05, -g 1.35e8. Details of enriched regions are available in Supplemental Data 4.

### ATAC-seq and data analyses

ATAC-seq was performed with two biological replicates. Nuclei were extracted from 1% formaldehyde-fixed seedlings, stained with 0.5 μM DAPI, and sorted with BD Influx™ cell sorter (BD Biosciences). For each replicate, 50,000 of sorted nuclei were collected in Galbraith buffer and centrifuged at 2000g at 4 °C for 5 min. The nuclei were resuspended with a 20 μl Tn5 transposase (Illumina) reaction. The transposed DNA was purified with MinElute PCR Purification Kit (Qiagen) and amplified with selected Nextera index oligos (Illumina). Size selection of PCR products was performed with AMPure® XP beads (Beckman Coulter) to collect library molecules between 200 and 700 bp. Finally, the purified libraries were pooled and sequenced. Raw reads were mapped to the TAIR10 *Arabidopsis* reference genome with Bowtie2 and sorted with SAMtools. The mapped reads were used to generate coverage files with 100 bp windows using bedtools (v2.26.0).

### Gene expression analyses

RNA sequencing (RNA-Seq) was performed with two biological replicates per sample. Total RNA was extracted from aerial parts of seedlings using a RNeasy Plant Mini Kit (Qiagen). Libraries were constructed with the NEBNext® Ultra™ RNA Library Prep Kit for Illumina (NEB E7770) according to manufacturer’s instructions. The RNA-Seq libraries were sequenced at Novogene (Cambridge, UK) via a Novaseq instrument in 150-bp paired-end mode. Reads were mapped to TAIR10 genome with HISAT 2 (v2.2.1) in paired-end mode. Differentially expressed genes (DEGs) were identified via the Subread and DESeq2 R packages with a cutoff of log2 fold change larger than 1.6 and false discovery rate (FDR) less than 0.01. Details of the reads count table and differentially expressed genes are available in Supplemental Data 2.

## Data Availability

Short read data of *in situ* Hi-C, ChIP-seq, ATAC-seq, and RNA-seq are publicly available at NCBI Sequence Read Archive under accession number PRJNA1043456.

Large datasets, such as normalized Hi-C matrices and BigWig ChIP-seq and ATAC-seq track files are available in the figshare repository. They are accessible with the following link: https://figshare.com/s/8a653d26c484048b67bf with Digital Object Identifier (DOI) 10.6084/m9.figshare.24533263. The epigenetic profile of various histone marks is available from our previous publication^21^.

## Code Availability

All scripts used in this study are available upon request.

## Acknowledgements

We thank Manuela Knauft and Jaqueline Pohl for their excellent technical assistance during Hi-C library preparation and sequencing. We thank Arp Schnittger for providing us *wapl1/2* mutants. We thank computing support by the High Performance and Cloud Computing Group at the Zentrum für Datenverarbeitung of the University of Tübingen, the state of Baden-Württemberg through bwHPC and the German Research Foundation (DFG) through grant no. INST 37/935-1 FUGG. This work was supported by DFG under grants No. JI347/6-1 and LI 2862/8-1, the Federal Ministry of Education and Research (BMBF) under grant No. 100489995, and the European Research Council (ERC) under the European Union’s Horizon 2020 research and innovation programme (grant agreement No. 757600).

